# *Zm*PMI3 phosphomannose isomerase kinetics is associated with natural variation of water-soluble carbohydrate content in mature maize stem

**DOI:** 10.64898/2026.10.12.758442

**Authors:** Stefan Robertz, Shaogan Wang, Frank Roeber, Susanne Groh, Markus Pauly, Vicente Ramirez

**Affiliations:** Institute for Plant Cell Biology and Biotechnology, Heinrich Heine University Düsseldorf, 40225 Düsseldorf, Germany; Bioeconomy Science Center (BioSC), Forschungszentrum Jülich, 53435 Jülich, Germany; Pioneer Hi-Bred Northern Europe Service Division GmbH, 79427 Eschbach, Germany

**Keywords:** *Zea mays*, water-soluble carbohydrates, maize stover, GWAS, phosphomannose isomerase, *Zm*PMI3, carbohydrate metabolism

## Abstract

Mature maize stems accumulate substantial amounts of water-soluble carbohydrates (WSC) providing an accessible carbon pool for industrial bioconversion. However, the genetic factors controlling WSC reserves under field conditions remain poorly resolved. In this study, we quantified total WSC including glucose, fructose, sucrose contents in the internode below the first ear of a genetically diverse maize panel at physiological maturity and associated trait variation with genome-wide SNP data. The panel displayed extensive variation in stem WSCs content, with values ranging up to 50.4 % of dry weight. Genome-wide association mapping identified 20 quantitative trait loci associated with individual sugars or total WSC content. One fructose associated region contained two adjacent gene annotations predicted to encode fragments of a phosphomannose isomerase. Sequence comparison, cDNA amplification and protein structure prediction supported a unified gene model, here designated as *ZmPMI3*. Biochemical characterization showed *Zm*PMI3 catalyzes the reversible conversion between mannose-6-phosphate and fructose-6-phosphate, a central reaction in carbohydrate metabolism. Natural protein haplotypes with distinct kinetic properties were identified correlating with significant differences in stem WSC content. Together, these findings identify *Zm*PMI3 as a candidate contributor to natural variation in maize stem carbohydrate reserves, potentially through altered hexose-phosphate partitioning.

## Introduction

Maize (*Zea mays* L.) is one of the world’s most important crops, yet its stover, comprising a substantial fraction of its vegetative biomass in the form of leaves and stems, is often underutilized after harvest (van der Weijde *et al*., 2013; Bichot *et al*., 2018; Soreng *et al*., 2022). In addition to structural carbohydrates embedded in lignocellulosic cell walls, mature corn stems contain large amounts of water-soluble carbohydrates (WSC) which are of particular interest because they represent a readily accessible carbon source for biotechnological applications (Gomez-Flores *et al*., 2018; Bautista, Unpaprom and Ramaraj, 2019; Robertz *et al*., 2024). The dominant WSC are sucrose (Suc) and its monosaccharide constituents glucose (Glc) and fructose (Fru) (Akter *et al*., 2020; Robertz *et al*., 2024). This soluble fraction is attractive for microbial bioconversion, because it is more accessible than recalcitrant lignocellulosic wall polysaccharides and can thus influence the efficiency with which stover is processed (Robertz *et al*., 2024).

Stem WSC accumulation is also biologically important. In maize, Suc is the primary product of photosynthesis, synthesized mainly in the mesophyll cells of source tissues from photosynthetically fixed CO_2_ (Seifert, 2004; Ruan, 2014). It is subsequently transported via the phloem to sink tissues, such as meristems, developing leaves, roots, flowers and seeds, where it is stored or split into Glc and Fru and further metabolized (Verbančič *et al*., 2018). The distribution and fate of these water-soluble carbohydrates is governed by dynamic source-sink relationships, i.e. the balance between source activity and sink demand (Slewinski, 2012). When photosynthetic carbon fixation exceeds immediate metabolic requirements, surplus WSC are stored (Stitt and Zeeman, 2012). In many plants, transient leaf starch acts as the dominant short term storage pool that fluctuates diurnally and supports nocturnal metabolism (Stitt and Zeeman, 2012; Miyake, 2016). Grasses, such as maize, sorghum, or sugar cane can also accumulate substantial amounts of WSC in the vacuoles of stem parenchyma cells, with reported levels ranging from 5 % to 35 % in mature maize plants (Chen *et al*., 2007; Bian *et al*., 2015; Cazetta and Revoredo, 2018; Akter *et al*., 2020).

Unlike the transient starch reserves in leaves, WSC pools accumulated in stem tissue function as longer-term storage (Slewinski, 2012). These reserves primarily sequester excess assimilates and therefore do not directly compete with other sink tissues under non-limiting conditions (Slewinski, 2012). During vegetative growth, expanding stems act as major carbohydrate sinks, whereas developing kernels become the dominant sink after the transition to the reproductive phase (Liang *et al*., 2019). This shift coincides with a decline in photosynthetic activity in source leaves, raising the possibility that stem carbohydrate reserves may contribute to grain filling, as observed in rice (Wang *et al*., 2017). However, studies in maize indicate that total stem WSC content remains relatively constant during the reproductive phase, suggesting that these reserves do not substantially contribute to seed filling (Liang *et al*., 2019). Instead, they act as a buffering system that can stabilize plant carbon metabolism during periods of environmental stress, such as drought, low light, or nutrient limitation, when photosynthetic carbon assimilation is reduced (Setter, Flannigan and Melkonian, 2001; Slewinski, 2012; Sekhon *et al*., 2016; Cazetta and Revoredo, 2018).

Previous studies have shown that stem sugar related traits in maize display moderate to high heritability, indicating that natural variation in WSC content is, at least in part, genetically controlled (Van Reen and Singleton, 1952; Slewinski, 2012; Bian *et al*., 2015). Less is known about the genetic architecture of individual WSC components in mature stem tissue across broad natural variation panels. A clearer understanding of this variation could identify loci useful for improving corn stover valorization and could reveal metabolic nodes that influence the accumulation of soluble carbohydrate reserves.

Here, we investigate the extent and genetic basis of natural variation in WSC accumulation in mature maize stems providing a framework for dissecting the genetic and metabolic regulation of stem WSC reserves in maize and highlight candidate entry points for future functional studies with relevance to crop resilience and crop residue valorization.

## Material and Methods

### Maize material

A total of 289 maize inbred lines used for the WSC profiling were grown in Grissheim, Baden-Württemberg, Planting date 14.5.2021, 10 plants per row, 1.4 m row length (2 m including alley). Sampling occurred 6 weeks after flowering. The inbreds were evaluated in an augmented block design including four genotypes (A619, B73, F7 and FAP1360A) as replicated checks. Stalk segments from 5 plants in the middle of the plot were sampled. At maturity, the internode immediately below the first ear was harvested, dried and used for subsequent analyses. The replicates of harvested samples are listed in Supplementary File 2.

### WSC quantification

The mature maize stalk was dried, milled (Wang *et al*., 2023) and the WSC were extracted and quantified as described (Robertz *et al*., 2024). In short, milled powder was resuspended in water and autoclaved for 15 min at 121°C using a VX-120 autoclave (Systec). Afterwards, the liquid fraction was filtered through PTFE membrane filters (0.2 µm) and used for water-soluble carbohydrate quantification on a high-performance anion-exchange chromatography (HPAEC) system equipped with a CarboPac PA20 column (3 x 150 mm) and a pulsed amperometric detector (PAD detector; Metrohm). The following gradient was used: 21 min at 1 mM NaOH, 9 min at 700 mM NaOH and 13 min at 1 mM NaOH, with a flow rate of 0.5 mL/min. The Glc, Fru and Suc contents were determined based on standards with known concentrations using Fucose as an internal standard.

### Genome-wide association study

Genome-wide association study (GWAS) was performed to identify genomic regions associated with variation in WSC traits in mature stalks. Best linear unbiased estimators (BLUEs) for each inbred line were calculated using the lme4 package (Bates *et al*., 2015) in R (version 4.1.2) with the following formula: lmer (Trait ∼ Genotype + (1|Entry:Plot)+(1|Replicate). Adjusted means for each inbred line were estimated using the *emmeans* package (Lenth and Piaskowski, 2026) and subsequently used as BLUEs for statistical analyses, correlation analyses and GWAS.

GWAS was performed using a mixed linear model (MLM) in which population structure was controlled by including the first two principal components derived from SNP data as fixed effects. A pairwise kinship matrix (K matrix) was calculated and incorporated as a random effect in the model to account for genetic relatedness among the 199 accessions and reduce spurious associations. Model fit was assessed using quantile-quantile (Q-Q) plots comparing observed and expected p-value distributions. Marker-trait associations were selected using a suggestive threshold of *p* < 1×10^-4^ used in similar studies before (Kaler *et al*., 2017; Nayyeripasand, Garoosi and Ahmadikhah, 2021; Panahabadi *et al*., 2021; Guo *et al*., 2023) and visualized using Manhattan plots. SNP positions were converted from the B73 RefGen_v2 to RefGen_v5 genome assembly using MaizeGDB (Woodhouse *et al*., 2021).

Quantitative trait loci (QTLs) were defined by extending upstream and downstream genomic region of each significant SNP according to chromosome-specific linkage disequilibrium (LD) decay (r^2^ = 0.1) (Supplementary Figure S3). Genes located within QTL intervals were extracted from EnsemblPlants (Yates *et al*., 2022) and annotated using MaizeMine (version 1.5). Candidate genes potentially involved in WSC metabolism were prioritized based on functional annotation.

### *ZmPMI3* gene model and haplotype-chemotype analysis

Genomic and cDNA sequences of *ZmPMI3* were obtained from MaizeGDB (Woodhouse *et al*., 2021), and the exon–intron structure was manually annotated based on these sequences. To investigate allelic variation in *ZmPMI3*, whole-genome sequencing data from a panel of more than 1,000 maize varieties was used (Grzybowski *et al*., 2023) of which a subset of 135 varieties overlapped with the diversity panel analyzed in this study. Non-synonymous SNPs within the coding sequence were identified and used to define haplotypes, with the B73 reference allele (RefGen_v5 genome) designated as haplotype 1. For each haplotype, mean values and standard deviations were calculated for Glc, Fru, Suc and total WSC content. The data were tested for normality and non-parametric tests (Kruskal–Wallis followed by Dunn’s test with Bonferroni correction) were applied where assumptions were violated, while normally distributed traits were analyzed using one-way ANOVA followed by Tukey’s HSD test (*p* < 0.05).

### *ZmPMI3* amplification

For gene amplification, B73 seedlings were grown in a greenhouse at 25 °C with a 16 h light/8 h dark photoperiod. Five-week-old stems were harvested and immediately ground with mortar and pestle to a fine powder in liquid nitrogen. The RNeasy extraction kit (Qiagen, cat. No. 74104) was used for total RNA extraction according to the manufactureŕs instructions. Samples were treated with TURBO™ DNase (ThermoFisher Scientific, cat. No. AM2238) for 30 min at 37°C to remove genomic DNA. GoScript™ Reverse Transcription Mix (Promega, cat. No. A2790) with Oligo(dT) primers was used for reverse transcription, according to the manufactureŕs instruction. The cDNA was then used to amplify and sequence *ZmPMI3* using gene specific primers (Supplementary table S1).

### Cloning of Zm*PMI3* haplotypes

The coding sequence of the *ZmPMI3* haplotype 1 (acc. Nr. EU974512.1) was synthesized (GeneArt gene synthesis, ThermoFisher Scientific) in a pDONR™221 backbone. *ZmPMI3* haplotype 2, 3 and 4 were generated by site-directed mutagenesis and Gibson assembly. Mutation-containing primers (Supplementary table S1) were designed with the NEBuilder tool, and overlapping fragments were amplified using Phusion™ high-fidelity polymerase. Purified PCR products were assembled according to the NEBuilder assembly tooĺs instructions and transformed into *E. coli* DH5α. Positive clones were selected by antibiotic resistance and confirmed by whole plasmid sequencing.

### *Zm*PMI3 expression, purification and *in vitro* activity

For prokaryotic expression of *ZmPMI3* haplotypes, the coding sequences of *ZmPMI3_hap1*, *ZmPMI3_hap2*, *ZmPMI3_hap3*, and *ZmPMI3_hap4* were amplified and cloned into the QIAexpress pQE-80L expression vector (Qiagen). The resulting vectors were transformed into *E. coli Rosetta™(DE3) pLysS* (Sigma-Aldrich) to express recombinant proteins. Protein expression was induced with 0.7 mM isopropyl-β-D-1-thiogalactopyranoside (IPTG) at 18 °C for 20 h. Cells were subsequently harvested and lysed by ultrasonication. Recombinant proteins carrying N-terminal 6×His tags were purified using a 1 mL HisTrap HP affinity column (GE Healthcare) connected to an ÄKTA purification system. The column was equilibrated with 5 column volumes (CV) of binding buffer (20 mM sodium phosphate, 0.5 M NaCl, and 20 mM imidazole, pH 7.4), followed by sample loading and washing with 30 CV of binding buffer. Bound proteins were subsequently eluted using a linear imidazole gradient from 20 to 250 mM. A total of 30 fractions (0.5 mL each) were collected during elution. The column was operated at a constant flow rate of 1 mL min⁻¹ throughout the purification. The presence and purity of the recombinant proteins were verified by SDS–PAGE using 8–16% Mini-PROTEAN® TGX Stain-Free™ protein gels (Bio-Rad). The concentration of purified proteins was determined using the Bradford reagent (Bio-Rad) based on a BSA standard curve.

The reaction from mannose-6-phosphate (M6P) to fructose-6-phosphate (F6P) was measured in a coupled enzymatic assay based on (Maruta *et al*., 2008) following the adaptations published in (Hu *et al*., 2016). This assay relies on the subsequent conversion from M6P via F6P to glucose-6-phosphate, which is then oxidized by a glucose-6-phosphate dehydrogenase to gluconate-6-phosphate, accompanied by the reduction of β-NADP^+^ to NADPH. All components were obtained from Sigma Aldrich. To each reaction purified protein (specific protein masses listed in Supplementary File 2) was added and mixed with substrate (M6P) in different concentrations ranging from 0 – 2800 µM. The applied molar extinction coefficient of NADPH corrected for the path length was 3.38 L/mmol. The Michaelis-Menten constants were calculated for four FPLC purified fractions (#18 -#21 Supplementary Figure S6) of each haplotype individually and used to calculate the average and standard deviation. Significant differences were identified with a One-Way ANOVA followed by Tukey’s HSD test (*p* < 0.05).

The reverse reaction from F6P to M6P was measured as an endpoint reaction by mixing 50 mM Tris-HCl buffer pH 7.5, 25 µL F6P (c = 2 mg/mL) and 228 ng of purified protein to a total volume of 300 µL. The samples were incubated with horizontal shaking at 400°rpm and 30°C for 10, 30, 40 and 60 min. Afterwards, the *Zm*PMI3 was inactivated by incubating for 10 min at 65°C. Each reaction was set up in duplicates. The resulting phosphate sugar mixtures were separated on a HPAEC system equipped with a CarboPac PA20 column (3 x 150 mm) and a PAD detector (Metrohm). The following gradient was used: 30 min 420 mM NaOH followed by flushing with 700 mM NaOH for 10 min and an equilibration phase of 15 min at 420 mM NaOH. The phosphate sugar concentration was then determined via a standard curve with known concentrations of M6P and F6P. Cellotetraose (Megazyme) was used as internal standard.

## Results

### A field-grown maize diversity panel reveals broad variation in stem WSC profiles

To quantify natural variation in mature stem carbohydrate reserves, we analyzed WSC content and composition in the internode below the first ear at maturity from a panel of 289 field-grown maize varieties (Figure 1 a, Supplementary File 2).

**Figure 1:**
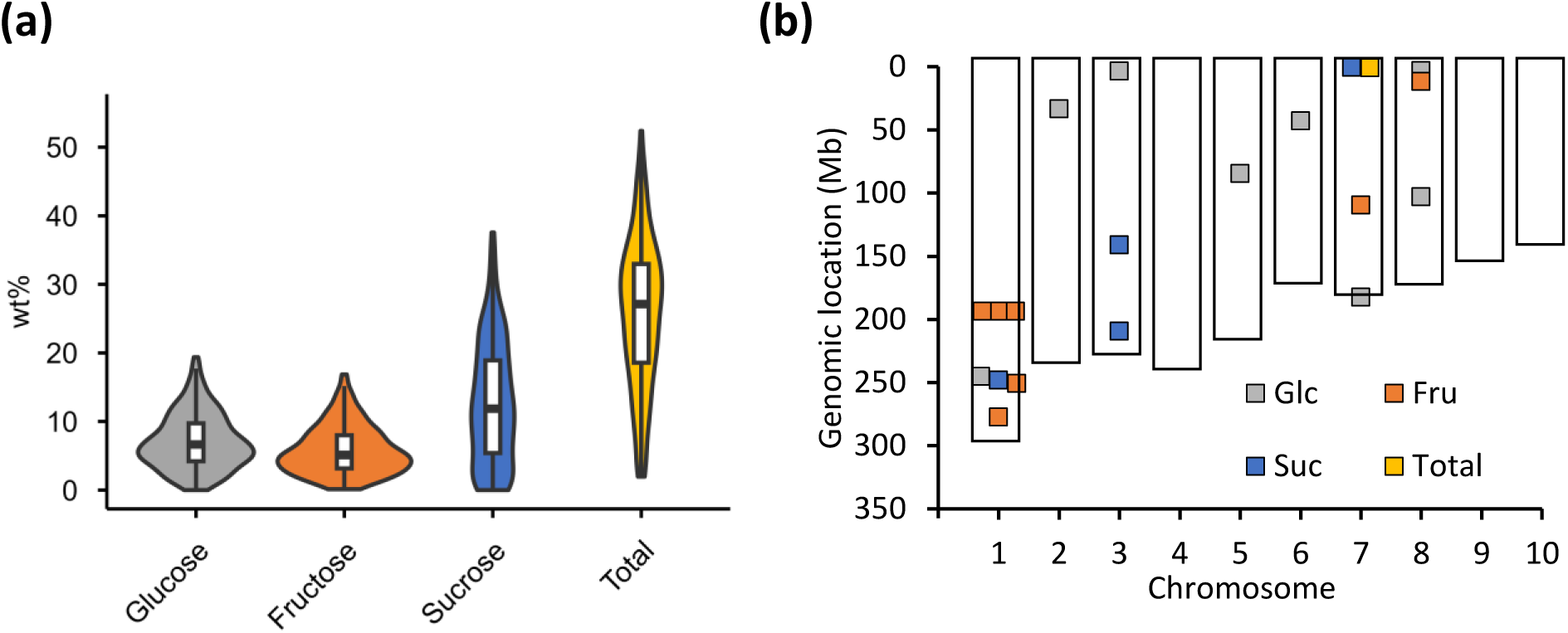
Water-soluble carbohydrate (WSC) content in corn stem and distribution of associated quantitative trait loci (QTLs) across the maize genome. **a).** The violin plots show the distribution of Glucose, Fructose, Sucrose and total water-soluble carbohydrate content (Total) in weight percent of total dry weight (wt%) across 289 different maize varieties. Boxplots within the violins indicate the median (horizontal line), interquartile range (upper and lower limit) and upper and lower whisker (vertical lines). Data are shown as the average per genotype with n = 3 for most genotypes. See Supplementary File 2 for details. b) Each data point represents one QTL detected for the different traits, i.e. the amount of glucose (Glc), fructose (Fru), sucrose (Suc) and the total water-soluble carbohydrate content (Total).

Substantial inter-genotypic variation was observed for Glc, Suc, Fru and total WSC content across the panel (Figure 1 a, Supplementary table S2). Suc was the predominant and most variable carbohydrate, with a mean content of 13.2 wt% and a range of 37.5 wt%. Several varieties, including Ky21 and Il14H, accumulated up to 37°wt% Suc, whereas others, such as SG18 or Hy contained only trace amounts. Glc and Fru were present at lower levels but also exhibited considerable variation. Glc content averages 7.1 wt% and ranged over 19.2 wt%, while Fru averaged 5.9 wt% with a range of 16.7 wt% (Figure 1 a, Supplementary table S2). Notably, varieties with high Glc content, such as Mo17 (18.5 wt%) and Hy (18.5 wt%), also accumulated high levels of Fru (16.4 wt% and 10.5 wt%, respectively), whereas their Suc content remained comparatively low (5.4 wt% in Mo17 and only traces in Hy). Total WSC content, calculated as the sum of Glc, Fru, and Suc, averaged 25.8 wt% across the panel and ranged by 51.2 wt% among genotypes (Figure 1 a, Supplementary table S2). In general, varieties with high Suc accumulation (Ky21 and IL14H) also exhibited the highest total WSC content.

### Genome-wide association study identifies multiple loci linked to mature stem WSC traits

To identify genomic regions associated with mature stem WSC content, best linear unbiased estimators (BLUEs) for Glc, Fru, Suc and total WSC were used for GWAS with a SNP matrix comprising 44,284 markers derived from genotyping with the Illumina MaizeSNP50 BeadChip across 199 maize varieties from the panel. Association mapping was performed using a mixed linear model that incorporated the first two genetic principal components and a kinship matrix to integrate by population structure and relatedness. Quantile-Quantile (Q-Q) plots confirmed the “goodness of fit” of the Mixed Linear Model, as the majority of observed *p*-values followed the expected distribution, while the significant associations deviated sharply from the expected *p-*values (Supplementary Figure S2).

Initially, 22 SNPs showed significant associations with one or more of the four traits analyzed, consistent with a polygenic architecture influenced by multiple small effect loci (Supplementary Figure S1 and Supplementary table S3). Two SNPs were excluded from further analysis due to inconsistencies between RefGen_v2 and the updated RefGen_v5 genome annotations (Supplementary table S3). For the remaining 20 SNPs, quantitative trait loci (QTLs) intervals were defined using chromosome-specific linkage disequilibrium (LD) decay (r^2^ = 0.1) (Supplementary table S4 and Supplementary Figure S3).

The 20 retained QTLs were distributed across 7 of the 10 maize chromosome (Figure 1 B). Eight QTLs were associated with Glc, seven with Fru, four with Suc, and one with total WSC content (Supplementary table S4). Furthermore, several Fru-associated QTLs on chromosome 1 overlapped (Supplementary table S4).

Manual inspection of genes annotated within the detected QTL intervals identified several candidates with possible roles in carbohydrate metabolism. Interestingly, the QTL Fru_8_1 contained two adjacent genes annotated as putative phosphomannose isomerases (PMIs) Zm00001eb335020 and Zm00001eb335030. Because PMI catalyzes the reversible interconversion between Fru-6-phosphate (F6P) and mannose-6-phosphate (M6P), a central reaction in carbohydrate metabolism, this locus was selected for further investigation.

### A Fructose-associated QTL contains a misannotated phosphomannose isomerase locus

Zm00001eb335020 (B73 RefGen_v5; chr8: 11.689.365 – 11.693.144) is predicted to encode a 168 amino acid protein, whereas Zm00001eb335030 (B73 RefGen_v5; chr8: 11.717.745 - 11.721.770) is predicted to encode a 244 amino acid protein located approximately 26 kb downstream. Both entries are flagged in MaizeGDB as potentially representing tandemly repeated genes with possible genome assembly/annotation issues. Inspection of the first coding sequence in the tandem revealed the absence of a stop codon, raising concerns about the proper annotation.

To further examine whether these two annotated genes represent independent genes or fragments of a single PMI, the predicted protein sequences were aligned with characterized PMIs from rice (*Os*PMI1, PMI2, PMI3) (Hu *et al*., 2016), Arabidopsis (*At*PMI1 and PMI2) (Maruta *et al*., 2008) and *Candida albicans* (*Ca*PMI) (Cleasby *et al*., 1996; Ahmad *et al*., 2018) (Figure 2 A - C; Supplementary Figure S4). The alignments revealed that the two maize protein sequences did not align with each other as simple duplicates. Instead, each sequence aligned to different portions of characterized PMI proteins (Supplementary Figure S4). The eukaryotic class 1 PMI consensus motif “YxDxNHKPE”, characteristic of zinc-dependent PMI enzymes (Coulin *et al*., 1993), was only present in Zm00001eb335020, together with three of the four zinc-coordinating residues previously identified in *Ca*PMI and conserved in *Os*PMI (Cleasby *et al*., 1996; Hu *et al*., 2016; Ahmad *et al*., 2018). The fourth was located in Zm00001eb335030 (Figure 2 B and Supplementary Figure S4). Furthermore, the predicted catalytic, C-terminal and helical domains defining *Ca*PMI (Cleasby *et al*., 1996; Ahmad *et al*., 2018) were split between the two maize protein sequences (Figure 2 B and Supplementary Figure S4). The predicted molecular weights of the two separated protein sequences were 18.5 kDa and 26.8 kDa, respectively, which are considerably lower than those reported for functional PMIs, such as *Os*PMI1 (46.9 kDa), *Os*PMI2 (44.2 kDa), *Chlorella variabilis* PMI1 (45.9 kDa) (Hu *et al*., 2016), *A. thaliana* PMI1 and PMI2 (48.5 kDa and 49.2 kDa, respectively) (Maruta *et al*., 2008) or *Ca*PMI (48.7 kDa) (Cleasby *et al*., 1996). In contrast, the combined molecular weight of both maize protein fragments is 45.3 kDa, consistent with full-length plant PMI enzymes.

**Figure 2:**
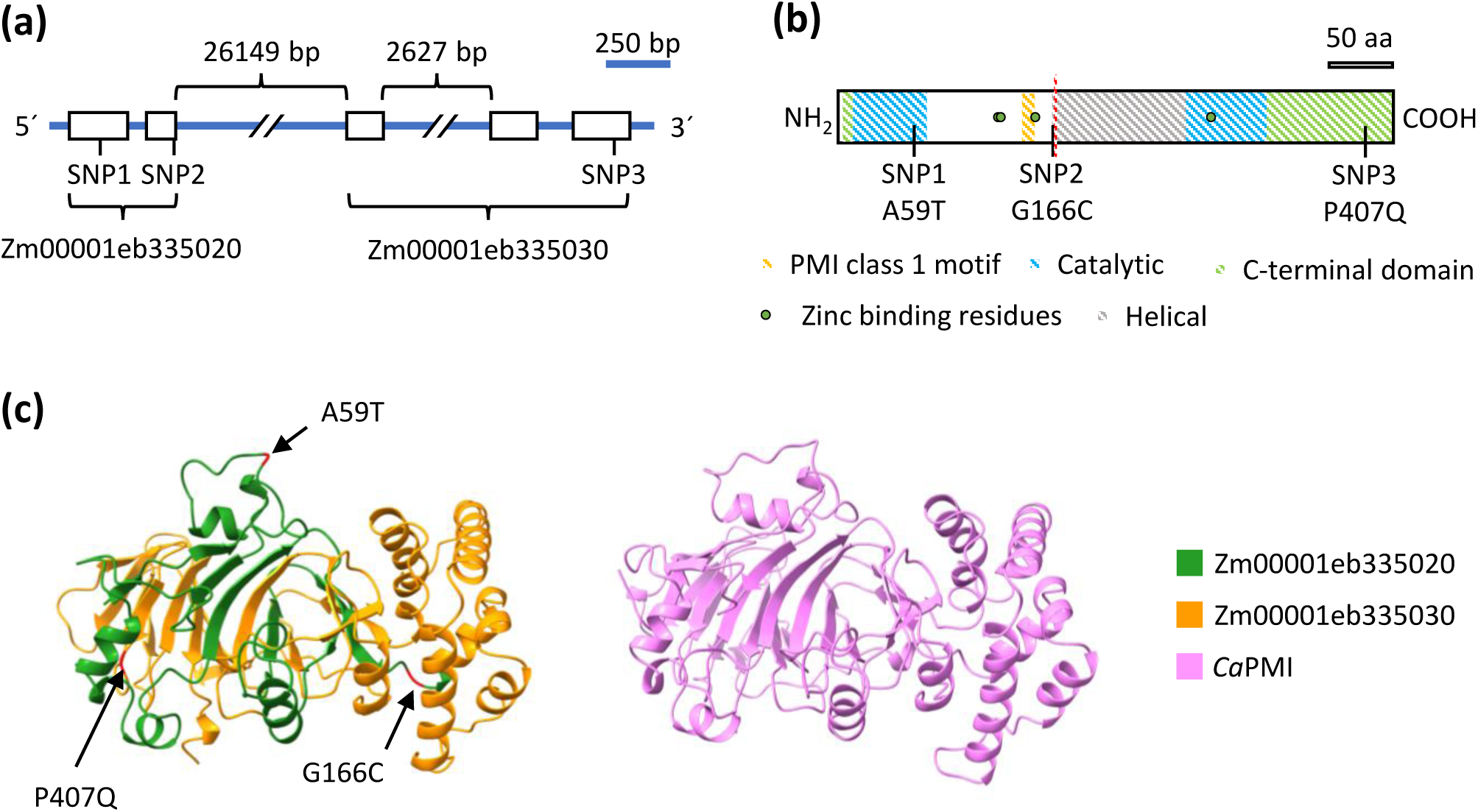
*PMI3* haplotype-chemotype analysis. **a**) Schematic representation of the SNP positions in the genomic region of *PMI3*. If not specified, the sizes of the exons (black boxes) and introns are acc. to the scale bar. **b**) Schematic representation of the SNP position in the PMI protein sequence. The PMI class 1 motif (orange), the predicted catalytic domains (blue), the helical domain (grey), the C-terminal domain (green), the Zinc binding residues (green dots) and the cut between protein sequence 1 and 2 (red) in the protein sequence of PMI3. Sizes are according to the scale bar. aa stands for amino acids. Domains are determined based on homology to *Ca*PMI and the results of (Ahmad et al., 2018). **c)** Structural comparison of ZmPMI3. Left: AlphaFold-predicted structure of *Zm*PMI3, with the regions corresponding to the two adjacent B73 RefGen_v5 PMI-like annotations shown in green (Zm00001eb335020) and orange (Zm00001eb335030). The positions of the three non-synonymous SNPs defining the ZmPMI3 haplotypes are indicated. Right: experimentally determined structure of *Candida albicans* PMI (CaPMI; PDB 5NW7), shown in pink, used as a structurally characterized class I PMI reference.

cDNA amplification from B73 stem tissue provided experimental evidence supporting the presence of a single transcript spanning the two annotated gene models (Supplementary Figure S5). PCR amplification from cDNA yielded a fragment encompassing portions of both annotated genes, demonstrating transcript continuity and indicating that they constitute a single open reading frame. Sanger sequencing of the amplicon confirmed its identity and matched the predicted coding sequence. In addition, mining of available full-length cDNA databases identified a maize sequence (NCBI accession Nr. EU974512.1) encompassing both genes sequences within a single open reading frame (Alexandrov *et al*., 2009; Soderlund *et al*., 2009). The resulting unified gene model, corresponding to LOC100272536, consists of five exons and four introns, including a large intron of approximately 26 kb (Figure 2 A), and encodes a predicted protein of 435 amino acids with a molecular weight of approximately 48 kDa.

To further assess whether the unified *ZmPMI3* gene model encodes a canonical phosphomannose isomerase, we predicted the three-dimensional structure of the corresponding protein using AlphaFold and compared it with a structurally characterized PMI from *Candida albicans* (Figure 2 C). The AlphaFold-predicted structure of *Zm*PMI3 resembled the canonical architecture of class I PMIs comprising a catalytic jelly-roll domain containing the conserved active-site and zinc-coordinating residues associated with PMI activity, together with helical and C-terminal domains that contribute to the overall enzyme architecture (Figure 2 C and Supplementary Figure S4). This structural organization supports the interpretation that the two adjacent maize annotations together encode a single class I PMI.

Since MaizeGDB currently annotates two PMI genes outside the Fru_8_1 region: *Zm*PMI1 (B73 RefGen_v5 Zm00001eb125530, chr3: 23.825.301 – 23.834.443) and *Zm*PMI2 (B73 RefGen_v5 Zm00001eb167360, chr4: 7.870.887 – 7.879.674), this unified locus is referred to as *ZmPMI3*.

### Natural *Zm*PMI3 haplotypes are associated with differences in total stem WSC content

To assess natural variation in *ZmPMI3*, we analyzed its coding sequence in 135 maize varieties included in the phenotyped panel for which whole-genome resequencing data were available (Grzybowski *et al*., 2023). Within the unified *ZmPMI3* coding sequence, three non-synonymous SNPs were detected and used to define four haplotypes (Figure 2 A/B and Figure 3 A). The most common haplotype, PMI3_hap1, was present in 85 varieties, including the reference variety B73. PMI3_hap2 was detected in 28 varieties and was characterized by SNP1 resulting in an A59T substitution in the predicted catalytic domain (Figure 2 B and Supplementary Figure S4). PMI3_hap3 was present in seven varieties and contained SNP2, resulting in a G166C substitution in the helical domain. PMI3_hap4 was present in six varieties and defined by SNP3, resulting in a P407Q substitution in the C-terminal domain (Figure 2 B and Supplementary Figure S4).

**Figure 3:**
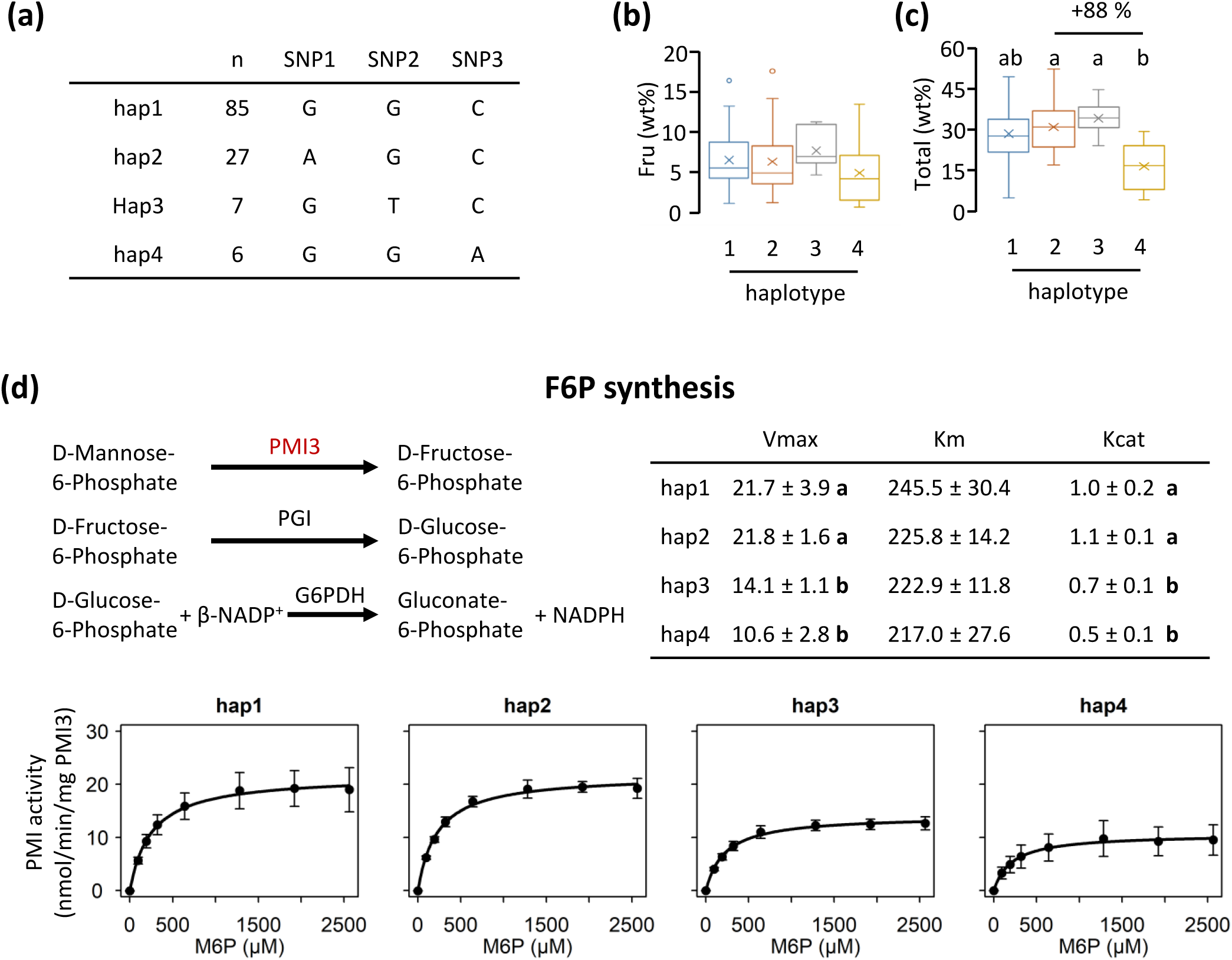
*PMI3* haplotype-chemotype analysis. **a**) Non-synonymous single nucleotide polymorphisms and their distribution across the individual haplotypes. n represents the number of varieties. **b**) Fructose (Fru) content and **c**) total water-soluble carbohydrate content (Total) in wt% (dry weight) of ZmPMI_hap1 to ZmPMI_hap4. Data are shown as the avg ± sd. Boxplots indicate the median (horizontal line), the average (cross), interquartile range (upper and lower limit), upper and lower whisker (vertical lines) and outliers (individual datapoints in b). No significant differences were found for Fru (b), determined with a Kruskal-Wallis test, followed by Dunńs multiple comparison test with Bonferroni adjusted p < 0.008, due to non-normal data distribution. Letters in c indicate statistically significant differences between the haplotypes, identified with a One-Way ANOVA followed by a Tukey-HSD test with p < 0.05. **d**) Fructose-6-phosphate (F6P) synthesis activity measured in a three-step enzyme cascade reaction. Kinetic parameters are displayed as Vmax (nmol/min/mg PMI3), Km (µM) and Kcat (1/s). The plots depict the Michaelis-Menten curve for each haplotype. Initial F6P synthesis activity was calculated based on co-factor formation with varied concentrations of M6P. All data are shown as the average ± SD of four different fractions from one protein purification experiment. Significant differences are indicated by the letters and were determined with a one-way ANOVA followed by a Tukey HSD/Kramer test with p < 0.05.

The three non-synonymous substitutions defining the *ZmPMI3* haplotypes neither map to the predicted active-site within the jelly-roll domain, nor to *Zm*PMI3 residues equivalent to other conserved *Ca*PMI zinc-coordinating residues defined in the characterized *Ca*PMI structure (Figure 2 C and Supplementary Figure S4). Alignment with functionally characterized and predicted plant PMIs further showed that the positions corresponding to SNP1 and SNP2 occur in relatively variable regions. In contrast, SNP3, which distinguishes PMI3_hap4 from PMI3_hap1, affects a position that is conserved among plant PMIs, including Arabidopsis PMI1 and PMI2, rice PMI1, PMI2 and PMI3, and the *Zm*PMI1 and *Zm*PMI2 putative maize paralogues, but not in fungal PMIs from *Candida albicans* or *Saccharomyces cerevisiae* or bacterial PMIs from *Escherichia coli* or *Salmonella typhimurium* (Wells *et al*., 1993; Cleasby *et al*., 1996; Maruta *et al*., 2008; Hu *et al*., 2016; Bangera *et al*., 2019) (Supplementary Figure S4). Because *ZmPMI3* was located with the QTL Fru_8_1, which is associated with the absolute Fru content in corn stover, Fru levels were compared among the four haplotypes (Figure 3 B). No statistically significant differences were detected among the four haplotypes (Figure 3 B). This lack of significance may reflect the smaller number of accessions available for haplotype analysis compared with the GWAS, the small size of rare haplotypes, or the possibility that the locus influences broader carbohydrate traits rather than Fru alone. In contrast, total WSC content differed among *Zm*PMI3 haplotypes (Figure 3 C). Lines carrying PMI3_hap2 and PMI3_hap3 showed significantly higher total WSC content than lines carrying PMI3_hap4 (Figure 3 C). Despite the small number of lines carrying haplotypes 3 and 4, these results are consistent with a possible involvement of PMI3 in the WSC metabolism in mature maize stems.

### *Zm*PMI3 encodes a functional PMI and natural haplotypes differ in enzyme kinetics

To test whether the unified *ZmPMI3* gene model encodes an active PMI, the four *Zm*PMI3 haplotypes were expressed in *E. coli* and purified by His-tag affinity chromatography. SDS-PAGE confirmed enrichment of recombinant proteins at the expected molecular weight (Supplementary Figure S6). PMI activity in the direction of F6P formation was measured using a coupled *in vitro* assay in which F6P produced from M6P was converted to G6P and subsequently oxidized with NADPH formation as the readout (Figure 3 D) (Maruta *et al*., 2008; Hu *et al*., 2016).

All four recombinant haplotypes displayed F6P synthesizing activity and followed Michaelis-Menten kinetics, indicating that PMI3 is a functional PMI (Figure 3 D). PMI3_hap1 and PMI3_hap2 exhibited similar kinetic parameters with V_max_ values of 21.7 and 21.8 nmol min^-1^ mg^-1^, respectively, and K_cat_ values of 1.0 and 1.1 s^-1^ (Figure 3 D). PMI3_hap3 and PMI3_hap4 showed decreased V_max_ values of 14.1 and 10.6 nmol min^-1^ mg^-1^, respectively, and decreased K_cat_ values of 0.7 s^-1^ and 0.5 s^-1^, respectively (Figure 3 D).

Compared to other plant-derived PMIs, *Zm*PMI3_hap1 exhibits lower catalytic efficiency than the enzymes reported from *A. thaliana* (Maruta *et al*., 2008). *At*PMI1 shows a higher affinity for M6P (K_m_ = 41.3 µM) and a higher velocity (V*max* = 1890 nmol/min/mg protein) (Maruta *et al*., 2008). In contrast, *At*PMI2 displays reduced substrate affinity (K_m_ = 372 µM) combined with a substantially higher velocity (V_max_ = 22500 nmol/min/mg protein) (Maruta *et al*., 2008). The two described rice PMIs exhibit higher catalytic rates than *Zm*PMI3 with V_max_ values of 24900 and 330 nmol/min/mg for *Os*PMI1 and *Os*PMi2, respectively (Hu *et al*., 2016). No enzymatic activity was detected for *Os*PMI3 (Os11g0600900) in (Hu *et al*., 2016). However, according to expression data from the Rice Expression Database (Xia *et al*., 2017) it is the most highly expressed PMI gene in rice. This discrepancy led to a closer inspection of *OsPMI3*. Notably, the NCBI protein entry appears to be misannotated, as it contains only a partial protein (227 amino acids), comparable to the misannotation of *Zm*PMI3. Further inspection of the rice genome identified an open reading frame at LOC4350861 encoding a 460 amino acid protein which likely represents the complete *Os*PMI3 protein, based on sequence alignment (Supplementary Figure S4). Additionally, PMIs from *Chlorella variabilis* and *Escherichia coli* showed substantially higher activities, with V_max_ values at 68000 and 74600 nmol/min/mg, respectively (Hu *et al*., 2016), which are in a range comparable to *Zm*PMI3. Since these proteins were purified with GST-tags, the K_m_ values cannot be compared, as the GST-tag likely influenced the M6P substrate affinity, as can be seen for the GST-tagged *At*PMI2 which exhibited a K_m_ value of 4104.7 µM (Hu *et al*., 2016).

The reverse reaction was assessed as an endpoint measurement by HPAEC-PAD quantification of M6P formation from F6P over a time course (Supplementary Figure S7). PMI3_hap2 produced more M6P from early timepoints than the rest of the haplotypes, apparently reaching a saturation stage after 40 min. PMI3_hap4 seems to produce slightly less M6P than PMI3_hap1 although only significant at timepoint 30 min (Supplementary Figure S7). These results support reversible PMI activity for *Zm*PMI3. Taken together, the combination of both *in vitro* activities confirms that PMI3 is a functional phosphomannose isomerase. In addition, SNP3, determining PMI3_hap4, seems to have a substantial impact on the enzyme activity, as both F6P and M6P synthesis activities are reduced. In combination, our results are consistent with a link between reduced *Zm*PMI3 activity associated with lower total WSC content.

## Discussion

### Natural variation in mature stem WSC profiles provides a basis for candidate-locus discovery

The accumulation of WSC reserves in maize stem tissue at physiological maturity has been recognized for decades (Singleton, 1948; Van Reen and Singleton, 1952). However, comprehensive analyses across large panels of field-grown maize varieties remain limited (Bian *et al*., 2015; Liang *et al*., 2019; Goñi *et al*., 2024). Stem WSC contents are strongly influenced by development, source-sink dynamics, and stem position (Slewinski, 2012; Sekhon *et al*., 2016; Liang *et al*., 2019; Akter *et al*., 2020). By consistently sampling the internode below the first ear at maturity, this study minimized these temporal and tissue-specific effects, enabling a more robust content comparison across a diverse maize panel.

The resulting dataset revealed a broad natural variation not only in total WSC content, but also in Glc, Fru and Suc composition. Suc represented the dominant and most variable component of the mature stem WSC pool, while Glc and Fru also varied substantially across the panel (Figure 1 A). The range of Suc variation observed was larger than that reported in some earlier studies based on smaller genotype sets or Brix measurements (Bian *et al*., 2015; Sekhon *et al*., 2016). This difference likely reflects both the larger genetic diversity captured here and differences in phenotyping methodology. Brix measurements estimate soluble solids in pressed stem juice, whereas the extraction approach used in this study quantified Glc, Fru and Suc from milled stem tissue and therefore captured a defined set of soluble carbohydrates from the sampled material. Differences in population structure, namely recombinant inbred lines versus a natural variation panel, may further contribute to the contrasting results. The range observed here is consistent with recent near-infrared spectroscopy estimates across large collections of maize samples (Goñi *et al*., 2024), supporting the conclusion that mature maize stover can display broad variation in soluble carbohydrate reserves.

The GWAS performed here was designed to identify QTLs associated with the natural genetic variation in mature stem WSC content and composition. Several aspects of the dataset require cautious interpretation. Phenotypic data were collected from a single field season and location, limiting the assessment of environmental stability and genotype-by-environment effects. In addition, the SNP dataset had moderate marker density and the association threshold used was relatively permissive. Despite these limitations, the Fru-associated Fru_8_1 locus illustrates this candidate-discovery value. Although the association signal alone does not establish causality, this region contained two adjacent annotations predicted to encode PMI-like protein fragments. Since PMIs interconvert fructose-6-phosphate and mannose-6-phosphate, this locus represented a biologically plausible entry point for investigating natural variation in carbohydrate metabolism. By extension, the remaining QTLs provide a useful framework for identifying additional genes potentially involved in stem carbohydrate metabolism, particularly when associated regions contain candidates with relevant predicted functions.

### *Zm*PMI3 resolves a gene model ambiguity and expands the maize PMI family

Detailed analysis of the Fru_8_1 locus indicates that the two adjacent PMI-like annotations in B73 RefGen_v5, Zm00001eb335020 and Zm00001eb335030, are unlikely to encode two independent functional PMIs. Instead, multiple lines of evidence support a single gene model encoding a 435 amino acid PMI, here designated as *Zm*PMI3. At the protein level, only the combined Zm00001eb335020–Zm00001eb335030 sequence contained the conserved PMI motif and domain architecture, showed a predicted three-dimensional fold consistent with characterized class I PMIs, and had a predicted molecular weight comparable to known plant PMIs (Figure 2 A–C; Supplementary Figure S4). This interpretation is further supported by previously reported full-length maize cDNA spanning both regions (Alexandrov *et al*., 2009; Soderlund *et al*., 2009) and by cDNA amplification from B73 stem tissue confirming a transcript that connects the two annotated regions (Supplementary Figure S5). Consistent with this corrected gene model, recombinant *Zm*PMI3 catalyzed both F6P and M6P formation *in vitro* (Figure 3 D and Supplementary Figure S7). Together, these data indicate that the Fru_8_1 locus contains a single functional PMI gene that is split into two adjacent annotations in the current maize reference genome.

Resolving this gene model is relevant beyond the specific GWAS locus because plant PMIs remain comparatively under-characterized despite their central metabolic position. For many years, PMIs were considered absent or difficult to detect in several higher plants, although functional enzymes have since been characterized in Arabidopsis and rice (Reed *et al*., 2001; Maruta *et al*., 2008; Stoykova and Stoeva-Popova, 2011; Hu *et al*., 2016). In Arabidopsis, PMI1 contributes to ascorbic acid biosynthesis through the Smirnoff–Wheeler pathway, whereas PMI2 has been linked to sugar starvation responses and possible cell wall polysaccharide turnover (Wheeler, Jones and Smirnoff, 1998; Maruta *et al*., 2008). In maize, the annotated PMI family has so far been limited mainly to *Zm*PMI1 and *Zm*PMI2, which have been described through genome annotation and transcriptomic studies, but whose encoded proteins have not been biochemically characterized in detail. The identification and functional characterization of *Zm*PMI3 therefore expands the maize PMI family and adds a biochemically validated member to the limited set of characterized plant PMIs.

### Natural *Zm*PMI3 variation links enzyme properties with stem WSC phenotypes

The strongest evidence linking *Zm*PMI3 to mature stem carbohydrate phenotypes comes from the convergence of natural sequence variation, enzyme properties and WSC accumulation. Four *Zm*PMI3 protein haplotypes were defined by three non-synonymous SNPs, and the corresponding recombinant proteins differed in catalytic behavior. Most notably, PMI3_hap4 displayed reduced F6P-forming activity and was associated with the lowest total WSC content among the haplotypes. This provides a direct biochemical connection between naturally occurring amino acid variation in *Zm*PMI3 and differences in mature stem carbohydrate reserves.

The metabolic position of PMI provides a plausible framework for interpreting this association. PMI reversibly interconverts F6P and M6P (Maruta *et al*., 2008), thereby linking the F6P/G6P pool associated with sucrose, glucose and fructose metabolism to mannose-derived pathways (Figure 4). M6P can be converted to mannose-1-phosphate and subsequently to GDP-mannose, which supplies the major L-galactose pathway of ascorbate biosynthesis and other mannose-dependent anabolic processes, including pathways contributing to cell wall polysaccharides and glycoproteins (Figure 4; (Smirnoff and Wheeler, 2000; Ruan, 2014; Fenech *et al*., 2018; Paciolla *et al*., 2019)).

**Figure 4:**
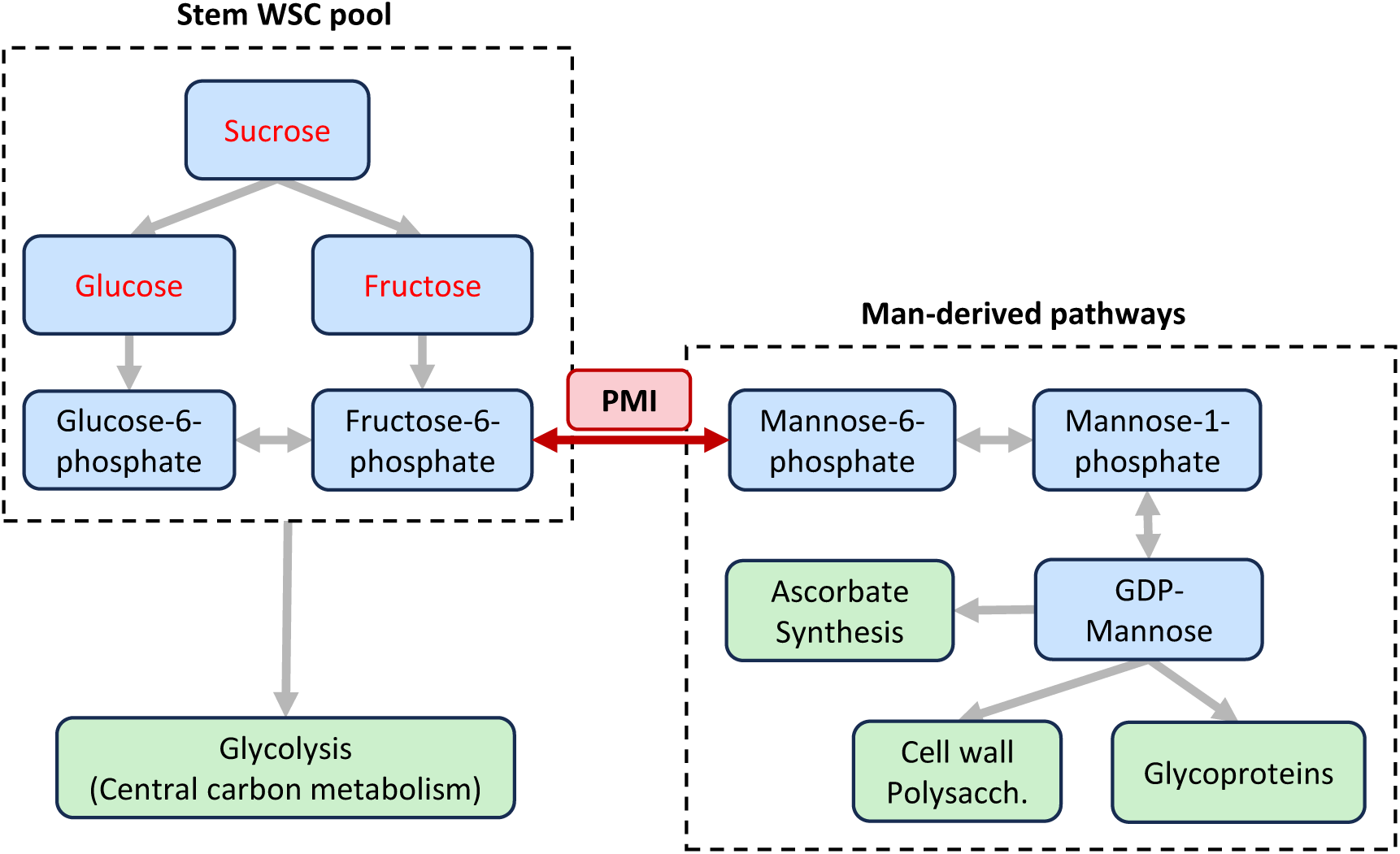
Metabolic context of *Zm*PMI3 in stem carbohydrate content. Phosphomannose isomerase (PMI) links the stem water-soluble carbohydrate (WSC) pool, which provides direct substrates for glycolysis, with mannose (Man)-derived metabolic pathways involved in ascorbate, cell wall polysaccharide (polysacch.), and glycoprotein biosynthesis. In this study, the stem WSC pool was quantified based on the abundance of sucrose, glucose, and fructose.

In mature maize stems, this branching point may be particularly relevant because WSC levels represent the balance between carbon retained as soluble reserves and carbon consumed, converted or remobilized through other metabolic pathways. Within this context, reduced F6P-forming activity of PMI3_hap4 could limit the return of carbon from M6P-linked metabolism into the F6P-associated central carbohydrate pool, thereby contributing to reduced accumulation or retention of WSCs during stem maturation. Such an effect would not require PMI to control WSC levels directly; rather, differences in PMI activity could alter one component of the broader metabolic network that determines how carbon is partitioned as the stem approaches physiological maturity.

Within this metabolic framework, the biochemical differences among *Zm*PMI3 protein haplotypes provide a possible mechanism linking natural variation at the Fru_8_1 locus with stem WSC composition. Changes in PMI activity could alter the balance between carbon retained within the F6P/G6P-linked central carbohydrate pool and carbon entering, or returning from, M6P-dependent metabolism. In principle, differences in this balance could propagate to sucrose, glucose and fructose pools through the extensive interconversion of soluble sugars and their phosphorylated intermediates. Accordingly, the reduced catalytic activity of some ZmPMI3 haplotypes could modify carbon partitioning at this branching point and thereby contribute to the WSC differences associated with the corresponding maize lines.

The relationship is unlikely to be simple or linear. PMI3_hap3 also showed reduced catalytic activity but was associated with higher total WSC content than PMI3_hap4. Net PMI flux *in planta* will depend not only on intrinsic enzyme kinetics, but also on substrate concentrations, reaction direction, protein abundance, tissue and developmental expression, environmental conditions and the demand imposed by downstream pathways. Nevertheless, the convergence of haplotype-specific enzyme properties with differences in mature stem WSC accumulation provides a functional link between natural ZmPMI3 variation and carbon partitioning, moving the interpretation of the Fru_8_1 locus beyond genetic association alone.

The structural context of the haplotype-defining substitutions is consistent with this interpretation. The substitutions are located outside the predicted active-site region (Figure 2 B) and do not affect *Zm*PMI3 residues equivalent to the catalytic Glu, other conserved *Ca*PMI residues required for isomerase activity, or the zinc-coordinating residues defined in the characterized fungal enzyme (Supplementary Figure S4). Accordingly, PMI3_hap4 retains activity in both reaction directions, indicating that the core isomerase mechanism remains intact. However, the SNP3-associated substitution distinguishing PMI3_hap4 from PMI3_hap1 occurs at a position conserved among the plant PMIs examined, but not among fungal or bacterial PMIs included in the comparison (Supplementary Figure S4). Although this conservation does not establish a specific functional role, it raises the possibility that this residue contributes to plant-specific tuning of PMI performance, for example through effects on local structural stability, domain positioning or conformational dynamics.

In conclusion, although the association between *Zm*PMI3 haplotypes and stem WSC traits remains to be functionally validated *in planta*, the combined evidence presented here supports *Zm*PMI3 as a compelling candidate linking natural genetic variation with mature stem carbohydrate reserves. These findings move beyond locus discovery by connecting GWAS, gene-model correction, and enzyme-level variation, thereby providing a foundation to investigate how central hexose-phosphate metabolism contributes to maize stover composition and carbohydrate-reserve regulation in grasses.

## Declarations

## Ethics approval and consent to participate

Not applicable

## Consent for publication

Not applicable

## CRediT authorship contribution statement

VR and MP conceptualized and supervised the work. SR and SW performed the experimental work. FR and SG designed the field trial and collected stalk sample. SR, VR, MP and SW wrote the manuscript. All authors approved the final version of the manuscript.

## Acknowledgements

The authors would like to thank Katharina Grosche for excellent technical support with the water-soluble carbohydrate screening.

---

**Supplementary Table S1:**
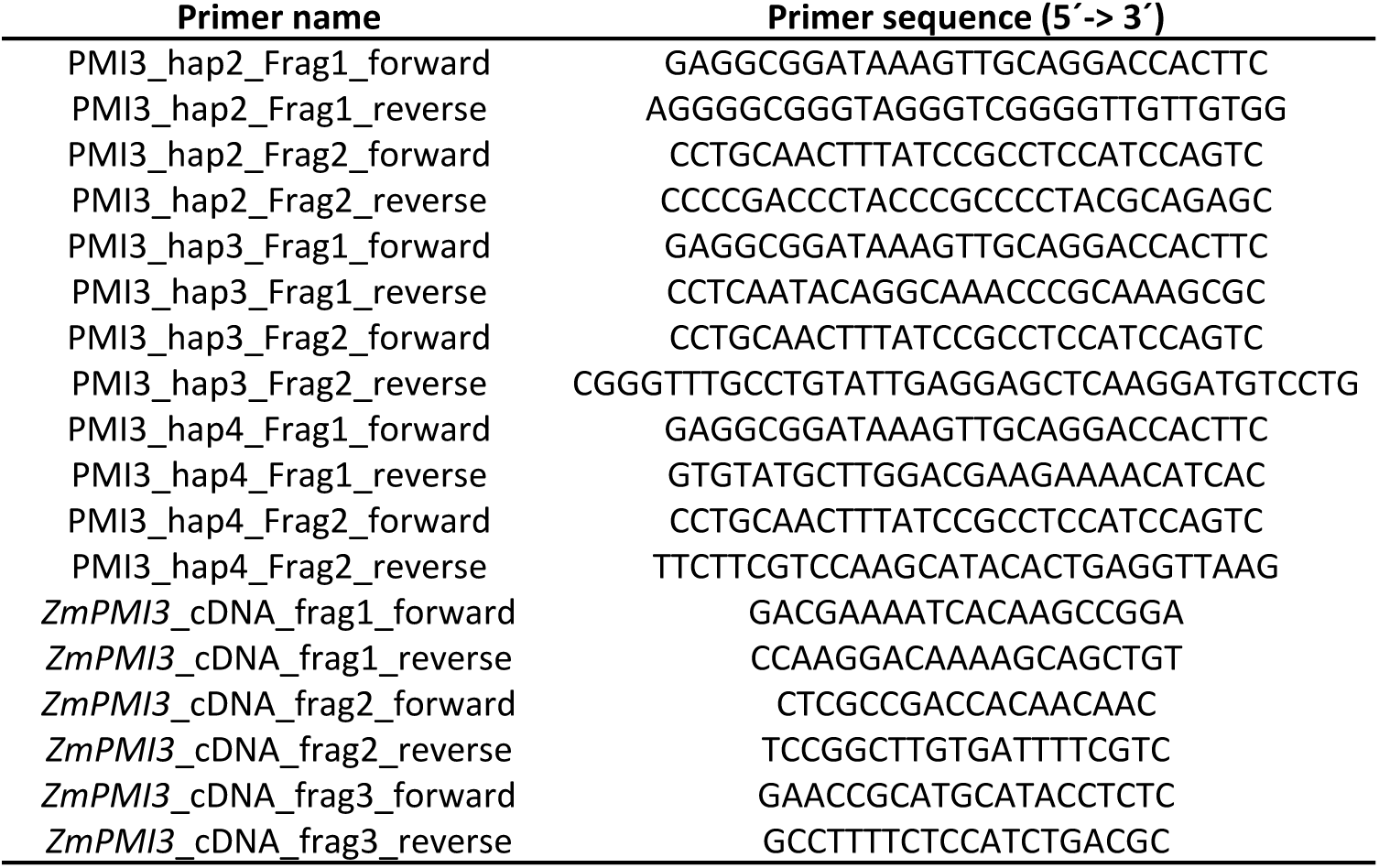
Primers used in this work.

**Supplementary Table S2:**
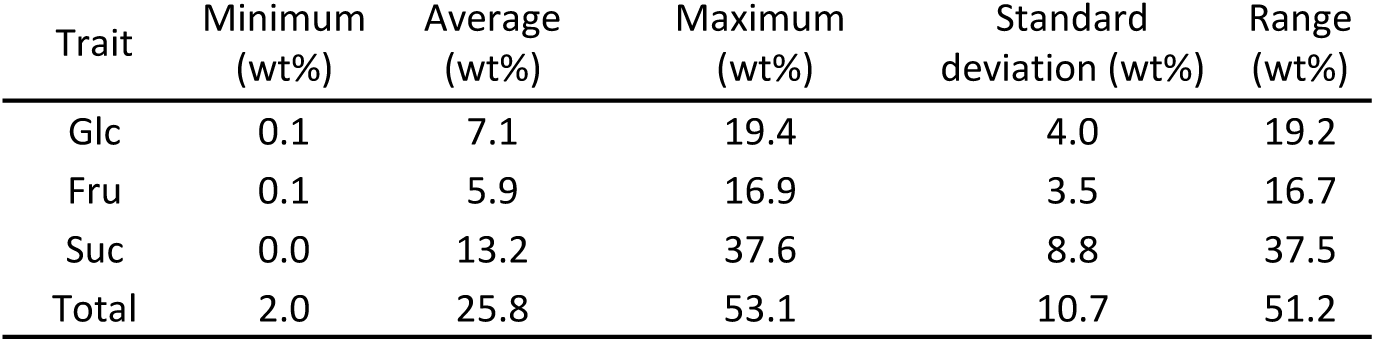
Descriptive statistics for the natural variation panel. Glucose (Glc), fructose (Fru), sucrose (Suc) and total water-soluble carbohydrate content (Total) in the natural variation panel.

| Trait | Minimum<br>(wt%) | Average<br>(wt%) | Maximum<br>(wt%) | Standard<br>deviation (wt%) | Range<br>(wt%) |
| --- | --- | --- | --- | --- | --- |
| Glc | 0.1 | 7.1 | 19.4 | 4.0 | 19.2 |
| Fru | 0.1 | 5.9 | 16.9 | 3.5 | 16.7 |
| Suc | 0.0 | 13.2 | 37.6 | 8.8 | 37.5 |
| Total | 2.0 | 25.8 | 53.1 | 10.7 | 51.2 |

**Supplementary Figure S1:**
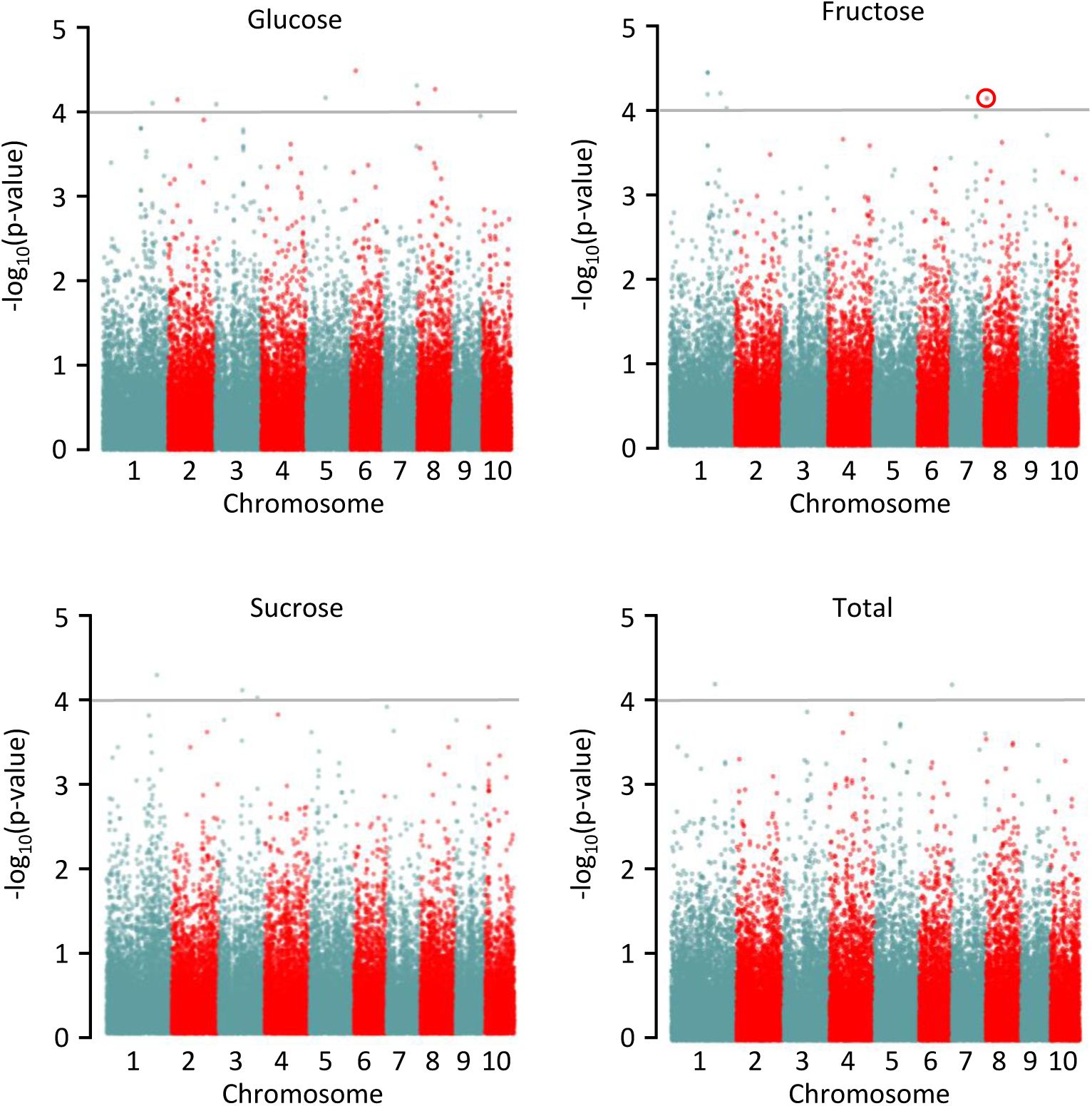
Manhattan-plots of genome-wide association study. The plots display the negative logarithm of the p-values for 44,293 SNPs across 199 corn varieties. Each dot represents one SNP, whose physical location in the genome is shown on the x-axis. Different colors distinguish the individual chromosomes. The horizontal line indicates the significance threshold of p < 0.0001. The traits are glucose (Glc), fructose (Fru), sucrose (Suc) and the total water-soluble carbohydrates (Total). Red circle indicates the SNP that defines the QTL which contains the *PMI3* gene.

**Supplementary Table S3:** Significant SNPs correlating to the 4 traits and their physical position in B73 RefGen_v2 and RefGen_v5. The analyzed traits are absolute content of glucose (Glc), fructose (Fru), sucrose (Suc) and total soluble carbohydrate content (Total).

| Trait | SNP ID/Locus | Chr | Physical position (bp) B73v2 | -log(P-value) | Synonyms |  | Physical position (bp) B73v5 |
| --- | --- | --- | --- | --- | --- | --- | --- |
| Glc | PZE.101192764 | 1 | 239535224 | 4.10 | rs131868092 | ss196431219 | 245178795 |
| Glc | SYN19476 | 2 | 31830162 | 4.14 | rs131181960 | ss196508254 | 33385853 |
| Glc | SYN6755 | 3 | 3701505 | 4.09 | rs131186087 | ss196526637 | 3755532 |
| Glc | PZE.105075111 | 5 | 82881987 | 4.16 | rs131538537 | ss196463853 | 84814765 |
| Glc | SYN26542 | 6 | 35091333 | 4.48 | rs130247068 | ss196513963 | 42964540 |
| Glc | PZE.107134586 | 7 | 173482344 | 4.31 | rs132449772 | ss196481431 | 182524830 |
| Glc | PZE.108002910 | 8 | 3159216 | 4.09 | rs131640778 | ss196481646 | 3467614 |
| Glc | PZE.108056460 | 8 | 101409730 | 4.26 | rs130826224 | ss196485451 | 103380217 |
| Fru | PZE.101196778 | 1 | 244906397 | 4.22 | rs128918254 | ss196431418 | 250699962 |
| Fru | PZE.101219678 | 1 | 271111712 | 4.04 | rs131892013 | ss196432348 | 277612674 |
| Fru | SYN28099 | 1 | 189586308 | 4.47 | No records found in MaizeGDB |  |  |
| Fru | SYN28100 | 1 | 189588905 | 4.47 | rs131826954 | ss196515234 | 193783734 |
| Fru | SYN28108 | 1 | 189589043 | 4.47 | rs131183513 | ss196515240 | 193783872 |
| Fru | SYN3135 | 1 | 189359106 | 4.21 | rs131826870 | ss196517981 | 193552471 |
| Fru | PZE.107053846 | 7 | 105221578 | 4.18 | rs130599888 | ss196478842 | 109896853 |
| Fru | PZE.108010908 | 8 | 11504308 | 4.16 | rs130713930 | ss196481916 | 11956275 |
| Suc | PZE.101195302 | 1 | 242541772 | 4.43 | rs128913628 | ss196431307 | 248216003 |
| Suc | PZE.103084194 | 3 | 139508679 | 4.24 | rs132034133 | ss196447370 | 141163822 |
| Suc | SYN8972 | 3 | 204363992 | 4.15 | rs129395180 | ss196528450 | 209675293 |
| Suc | PZE.107000798 | 7 | 988177 | 4.03 | rs131179431 | ss196475190 | 845208 |
| Total | PZE.101171014 | 1 | 214648677 | 4.26 | No records found in MaizeGDB |  |  |
| Total | PZE.107000798 | 7 | 988177 | 4.25 | rs131179431 | ss196475190 | 845208 |

**Supplementary Figure S2:**
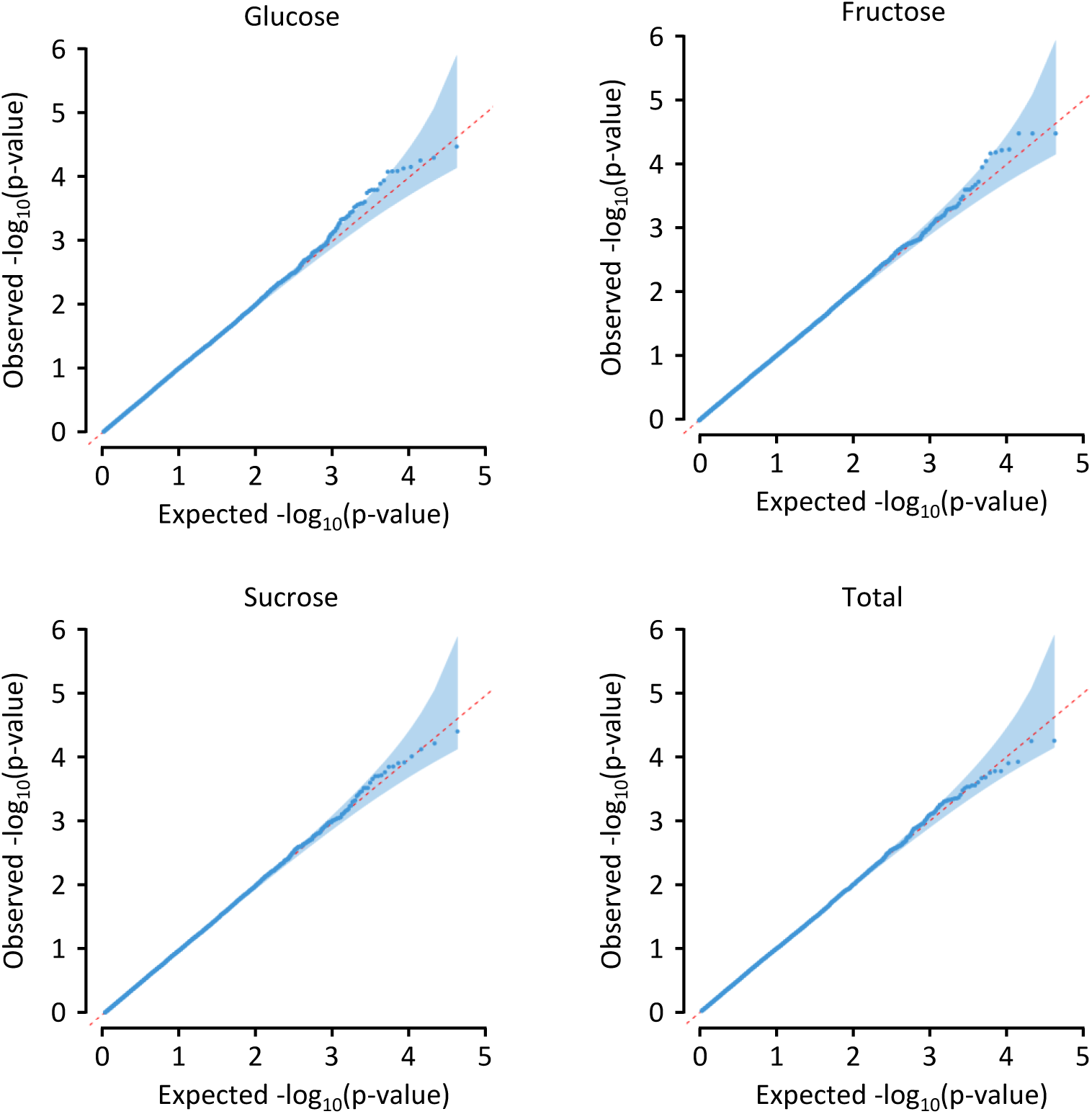
Quantile-quantile (Q-Q) plot of GWAS based on the mixed linear model for the traits Glucose, Fructose, Sucrose and total water-soluble carbohydrates. The x-axis and y-axis represent the expected −log_10_(p-values) and observed −log_10_(p-values), respectively. The dots indicate −log10(p) of the SNPs, the red line represents the expected values and the shaded area the 95 % confidence interval under the null hypothesis for no association.

**Supplementary Figure S3:**
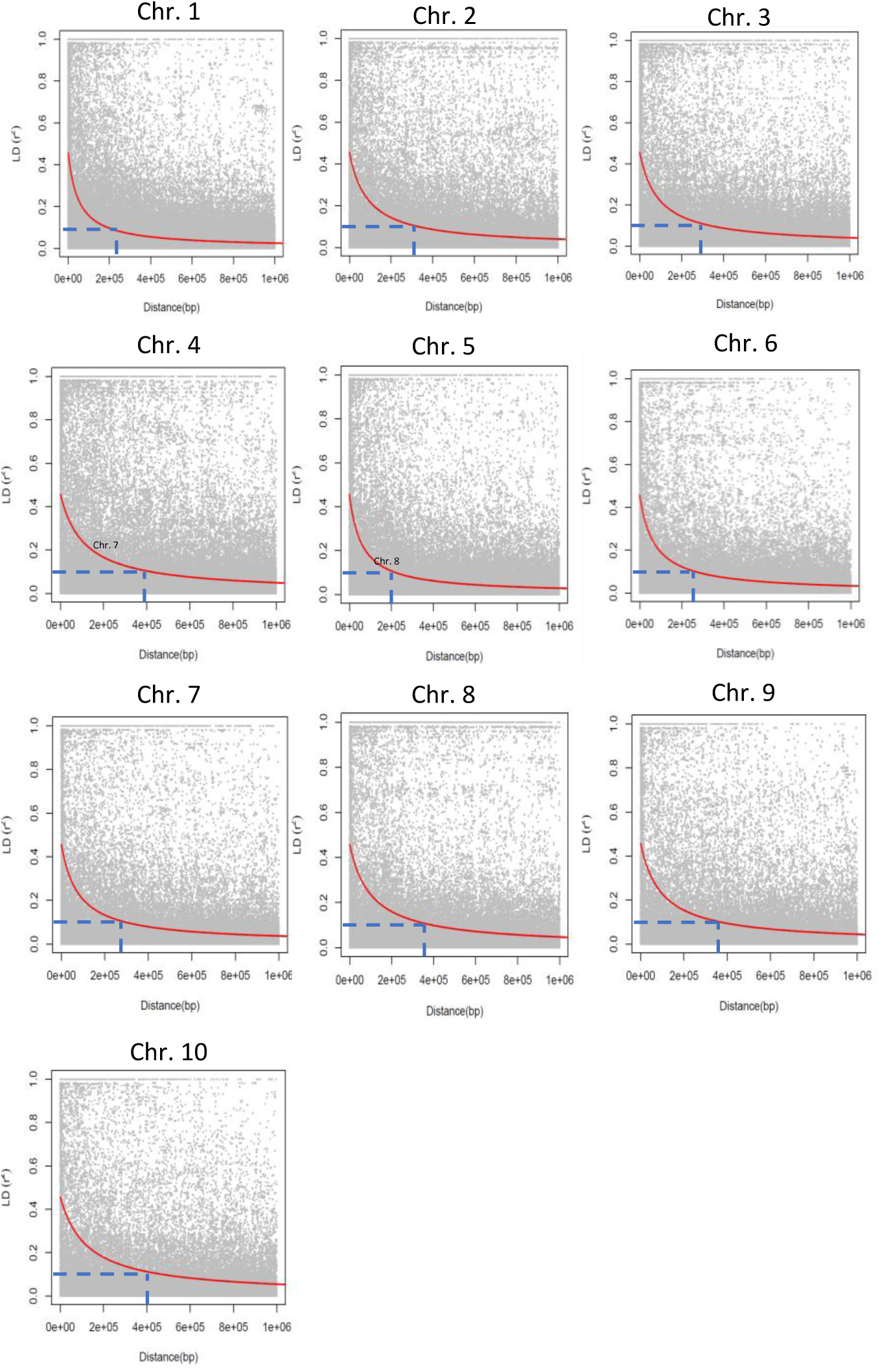
Chromosome specific linkage disequilibrium decay plots. Plotted is the r^2^ between the SNPs against the physical distance between them. The red line shows the linkage disequilibrium decay, and the blue bars show the threshold of 0.1 and its corresponding physical distance.

**Supplementary Table S4:** Overview of all QTL regions for water-soluble carbohydrate traits and candidate genes in maize. The traits are: the amount of glucose (Glc), fructose (Fru), sucrose (Suc) and the total water-soluble carbohydrate content (Total); * QTL Fru_8_1 contains candidate gene *PMI3*; Bold: the QTL Glc_6_1 is located in bin locus 6.01, previously associated with maize stalk Suc content (Bian et al., 2015), italics: the QTLs Fru1_1 to Fru1_3 overlap.

| QTL Name | Trait | Chr. | Physical position B73<br>RefGen_v5 |  | Predicted<br>genes in QTL |
| --- | --- | --- | --- | --- | --- |
|  |  |  | 5' start | Size |  |
| Glc_1_1 | Glc | 1 | 244978795 | 400 Kb | 9 |
| Glc_2_1 | Glc | 2 | 33085853 | 600 Kb | 13 |
| Glc_3_1 | Glc | 3 | 3455532 | 600 Kb | 32 |
| Glc_5_1 | Glc | 5 | 84614765 | 400 Kb | 6 |
| **Glc_6_1** | Glc | 6 | 42714540 | 500 Kb | 8 |
| Glc_7_1 | Glc | 7 | 182224830 | 600 Kb | 29 |
| Glc_8_1 | Glc | 8 | 3067614 | 800 Kb | 20 |
| Glc_8_2 | Glc | 8 | 102980217 | 800 Kb | 17 |
| Suc_1_1 | Suc | 1 | 248016003 | 400 Kb | 9 |
| Suc_3_1 | Suc | 3 | 140863822 | 600 Kb | 9 |
| Suc_3_2 | Suc | 3 | 209375293 | 600 Kb | 25 |
| Suc_7_1 | Suc | 7 | 545208 | 600 Kb | 16 |
| *Fru_1_1* | Fru | 1 | 193352471 | 400 Kb | 16 |
| *Fru_1_2* | Fru | 1 | 193583734 | 400 Kb | 16 |
| *Fru_1_3* | Fru | 1 | 193583872 | 400 Kb | 16 |
| Fru_1_4 | Fru | 1 | 250499962 | 400 Kb | 8 |
| Fru_1_5 | Fru | 1 | 277412674 | 400 Kb | 8 |
| Fru_7_1 | Fru | 7 | 109596853 | 600 Kb | 9 |
| Fru_8_1* | Fru | 8 | 11556275 | 800 Kb | 13 |
| Total_7_1 | Total | 7 | 645208 | 600 Kb | 12 |

**Figure S4:**
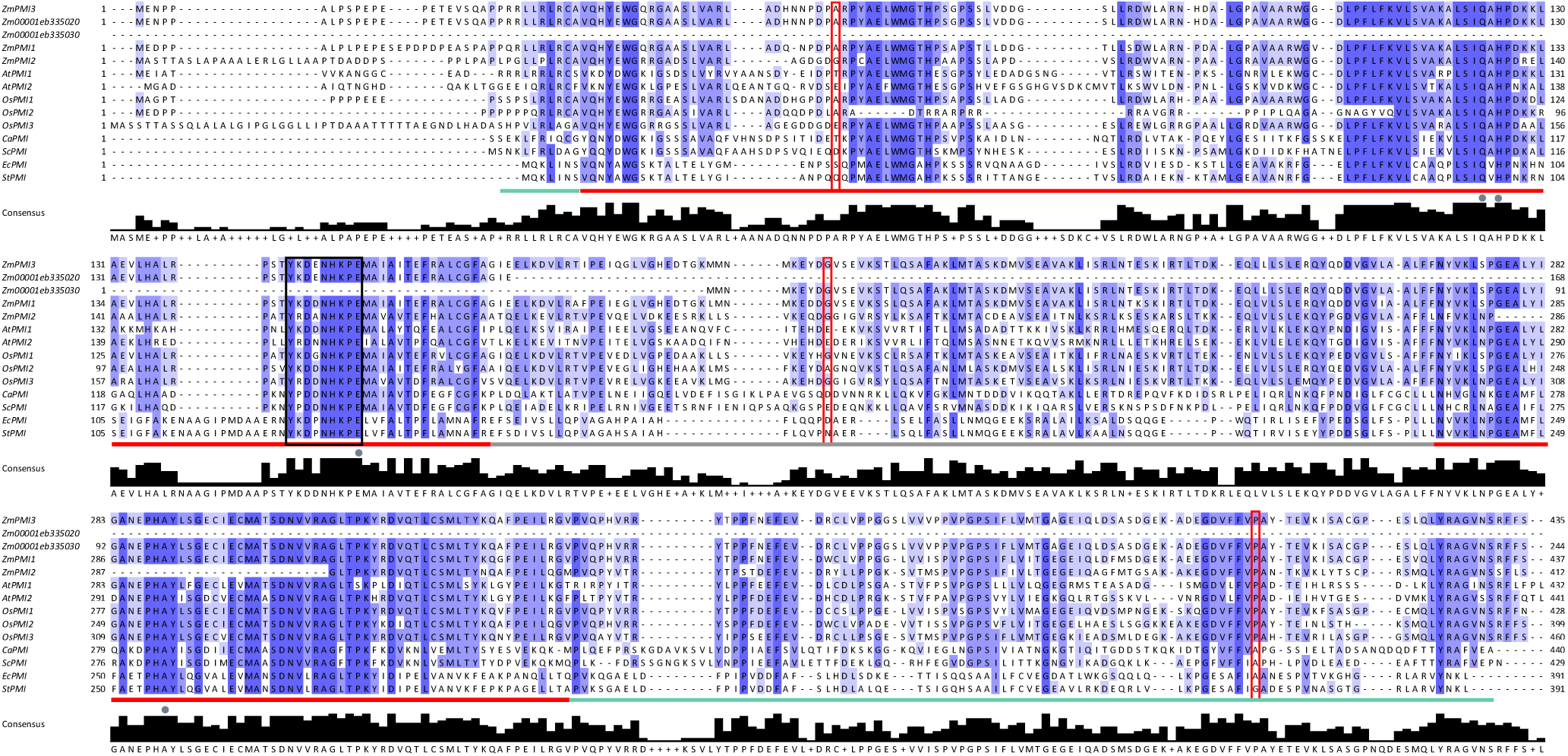
Protein sequence alignment of described and putative PMIs. The alignment was done with Clustal Omega (1.2.4) using the default parameters and visualized with Jalview (2.11.5.2). Identical and conserved amino acids across sequences are color coded in dark and light blue, respectively. Four zinc-ligand binding residues (grey dots; based on *Os*PMI1) and the PMI class 1 consensus sequence (black box) are indicated acc. to (Hu et al., 2016). Red boxes indicate the positions of the non-synonymous SNPs in *Zm*PMI3. The bars below the sequences indicate the catalytic domain (red), the helical domain (grey) and the C-terminal domain (green) based on *Ca*PMI (Ahmad et al., 2018). The histograms shows the degree of consensus with the most conserved amino acid listed below. The protein sequences were retrieved from MaizeGDB, NCBI or UniProt under the following accession numbers: *Zm*PMI1 Zm00001eb125530_T002; *Zm*PMI2 Zm00001eb167360_T001; *At*PMI1 Q9M884; *At*PMI2 Q9FZH5; *Os*PMI1 Q9FTR6 (Os01g0127900); *Os*PMI2 Os09g0389000; *Os*PMI3 Os11g0600900; *Ca*PMI (P34948); *Sc*PMI = *Saccharomyces cerevisiae* PMI (P29952); *Ec*PMI = *Escherichia coli* PMI (P00946); *St* = *Salmonella typhimurium* PMI (P25081).

**Figure S5:**
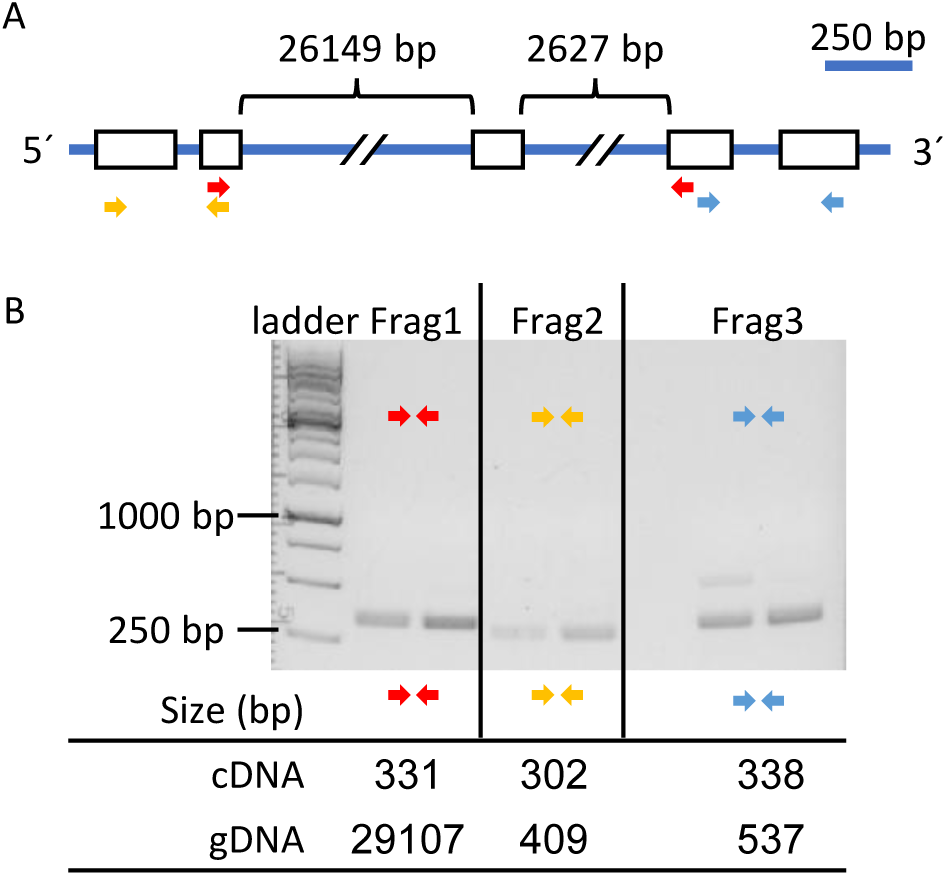
**A)** Schematic representation of the genomic region of *PMI3*. If not specified, the sizes of the exons (Ex) and introns are acc. to the scale bar. Colored arrows represent the position of the primers used to amplify the *PMI3* cDNA sequence. **B)** Individual fragments (Frag1 – 3) of *PMI3* were amplified from two individual B73 stem cDNA preparations and separated via gel electrophoresis. Colored arrows correspond to the arrows in A. Expected size refer to the amplicon size in complementary DNA (cDNA) and genomic DNA (gDNA). The sequences were confirmed via Sanger sequencing. Primer sequences are given in Supplementary Table 1. “Ladder” refers to the GeneRuler 1kb DNA ladder (Thermo Scientific).

**Figure S6:**
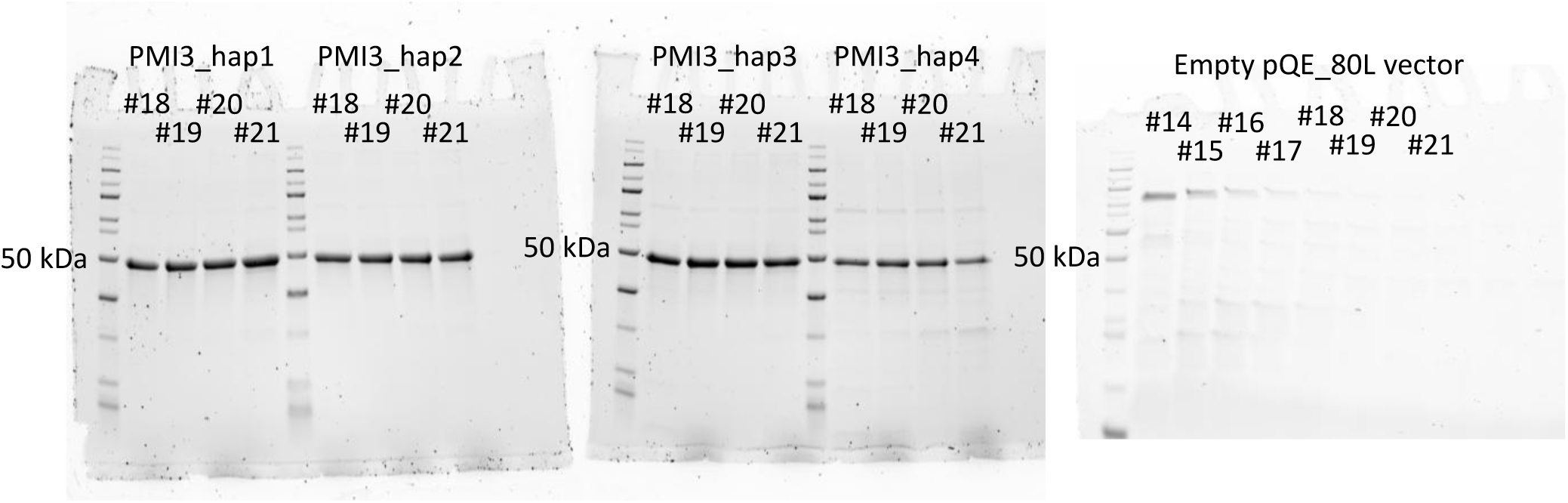
SDS-PAGE analyses of FPLC purified fractions of the distinct PMI3 haplotypes and the empty pQE_80L vector control. Of each fraction, 2.28 µg of total protein were added. Predicted molecular weight of ZmPMI3 = 47.7 kDa. Numbers refer to the individual fractions.

**Figure S7:**
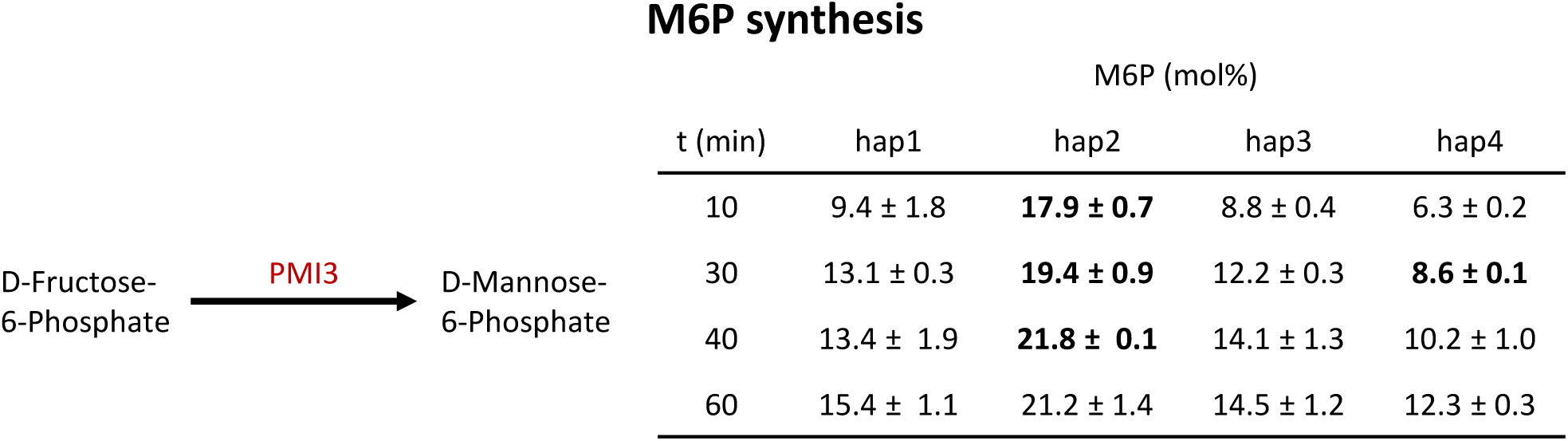
PMI activity of distinct PMI3 haplotypes. Mannose-6-phosphate synthesis (in mol%) measured over time (min) of the distinct PMI3 haplotypes. Data are shown as the average ± standard deviation of 2 independent reactions using the same purified protein. Bold values indicate significant different values compared to hap1 for each time point. Values were compared to haplotype 1 using two-tailed Students T-Test with p < 0.05.

